# Development of MHC Class I Blocking Peptides to Target Metabolic Dysfunction-Associated Steatohepatitis CD8^+^ T Cell Activation

**DOI:** 10.1101/2024.03.12.584666

**Authors:** Victoria Adams, Sudeep Sarma, Carol K. Hall, Arion Kennedy

## Abstract

MHC class I molecules play a crucial role in the immune system by presenting peptides derived from intracellular proteins to cytotoxic T lymphocytes (CTLs). This process is essential for immune surveillance and eliminating infected or malignant cells. In some diseases, the immune system fails to recognize and eliminate abnormal cells, leading to disease progression. Under conditions of metabolic dysfunction-associated steatohepatitis (MASH), subsets of CD8^+^ T cells have been identified as pathogenic, leading to inflammation and fibrosis. Therefore, explicitly targeting factors responsible for T cell activation may be necessary to prevent the onset of MASH and future complications such as cirrhosis or hepatocellular carcinoma. We have identified a specific MHC class I antigen that activates hepatic and splenic CD8^+^ T cells isolated from MASH mice. To specifically target the antigen, we developed two MHC H2-K^b^ blocking peptides, MHCP3 and MHCP5, that competitively inhibit the Ncf2 peptide from binding to H2-K^b^ and reduce activation and proliferation of CD8^+^ T cells. By inhibiting the recognition of specific antigens, these blocking peptides may prevent the activation of CD8^+^ T cells and progression of MASH.

## INTRODUCTION

Metabolic associated fatty liver disease (MAFLD) is one of the most common liver diseases. MAFLD represents a spectrum of liver pathologies that can transition from simple hepatic steatosis to MASH. This transition is a significant concern as MASH is characterized further by increased inflammation, fibrosis, and CD8^+^ T cell activation, which can ultimately develop into hepatocellular carcinoma. With no current FDA-approved treatments for MAFLD-related liver injuries, developing targeted therapeutic techniques is essential for preventing and treating this disease.

Previous work demonstrates that CD8^+^ T cells drive hepatic inflammation and fibrosis in mice and humans[1–5]. Recent work has identified multiple mechanisms that activate CD8^+^ T cells under MASH conditions. Metabolites, such as adenosine triphosphate and acetate, have been implicated in the development of auto aggressive hepatic CD8^+^ T cells[6]. In addition, the cytokine IL-15 is associated with activation in mouse models of MASH[7]. Classically, antigens presented by the major histocompatibility complex (MHC) class I activate CD8^+^ T cells. These antigens are peptides 8 to 9 amino acids in length and are collectively known as the MHC I immunopeptidome. The immunopeptidome is complex, contains both self and foreign immunogenic peptides, and is influenced heavily by protein translation and degradation. Metabolic diseases and infections also can alter the immunopeptidome. Antigens have been identified in diabetes, obesity, and other metabolic disorders and are essential in activating CD8^+^ T cells. Recently, Adams et al. 2023 identified a unique H2-K^b^ immunopeptidome in MASH conditions that contains the peptide Ncf2 (sequence: VHYKYTVV), which activates both hepatic and splenic MASH CD8^+^ T cells *in vitro*[1]. This suggests that blocking H2-K^b^ Ncf2 antigen presentation could provide a potential therapeutic benefit for treating MASH.

Blocking peptides have potential as a valuable tool for treatment of chronic inflammatory and autoimmune diseases[8, 9]. Blocking peptides are short amino acid sequences that, due to the small size, can interfere with protein-protein interactions. These peptides are emerging as valuable molecules in the drug discovery and development pipeline because of the high potency, low toxicity, and modest manufacturing costs. Given the identification of key peptides that activate MASH associated CD8^+^ T cells, designing blocking peptides that disrupt MHC class I dependent antigen presentation to CD8^+^ T cells may be an effective therapy for MASH.

Blocking peptides can prevent T cell activation either by binding to the T cell receptor (TCR) to prevent its association with a specific antigen or by binding to the MHC class I receptor to prevent T cells from interacting with specific antigens. There are several different ways to design blocking peptides. One common approach is to use sequence-based methods to predict binding of peptides with a high affinity for the MHC class I molecule of interest. Specific residues can be identified that are important for the activity of a targeted protein. An alternative to this approach is to computationally design peptides that bind specifically to the antigen and, thereby, block its activity. Here, we use the *Pep*tide *B*inding *D*esign algorithm (PepBD), a Monte-Carlo search algorithm in peptide sequence and conformation space, to discover peptides that bind with higher affinity and selectivity to a biomolecular target than a known “reference ligand.” The performance of the best peptide binders predicted by PepBD is evaluated *in-silico* by performing atomistic molecular dynamics (MD) simulations and calculating the binding free energy (ΔG_binding_) of the peptide:protein complex using the implicit-solvent molecular mechanics/generalized Born surface area (MM/GBSA) approach with the variable internal dielectric constant model[10–15]. Using the PepBD algorithm, we developed blocking peptides that competitively inhibit the interaction between Ncf2 peptide and H2-K^b^ to prevent CD8^+^ T cell activation and that might ultimately serve as an effective anti-MASH therapeutic strategy.

## RESULTS

### H2-K^b^ is a therapeutic target for MASH

We examined the role of CD8^+^ T cells in the development of MASH using low density lipoprotein receptor knockout (LDLRKO) mice on chow (control) or western diet (WD; MASH) for 12 weeks. The LDLRKO mice on WD represent an obese hyperlipidemic model of MASH and display a mild inflammation, fibrosis, and increased hepatic CD8^+^ T cells[3]. After 12 weeks, livers from control and MASH mice were collected to examine changes in the H2-K^b^ immunopeptidome. By using H2-K^b^ immunoaffinity chromatography and liquid chromatography with tandem mass spectrometry (LC-MS/MS), we identified antigens from livers of normal and MASH mice (Figure 1A). Candidate antigens were selected by searching the mouse proteome. The antigen datasets for all groups displayed the characteristic 8 to 11 amino acid length distribution for H2-K^b^ restricted antigens. BioVenn filtering analysis revealed 59 unique chow (control), 227 unique MASH, and 219 shared peptides. Of these peptides the Ncf2 (sequence: VHYKYTVV) and Gpnmb (sequence: FVYVFHTL) peptides were only identified in the MASH mouse model. NetMHCPan-4.1 was used to predict H2-K^b^ binding affinities and revealed that both the Ncf2 and Gpnmb peptides were strong binders (Figure 1B).

**Figure 1.**
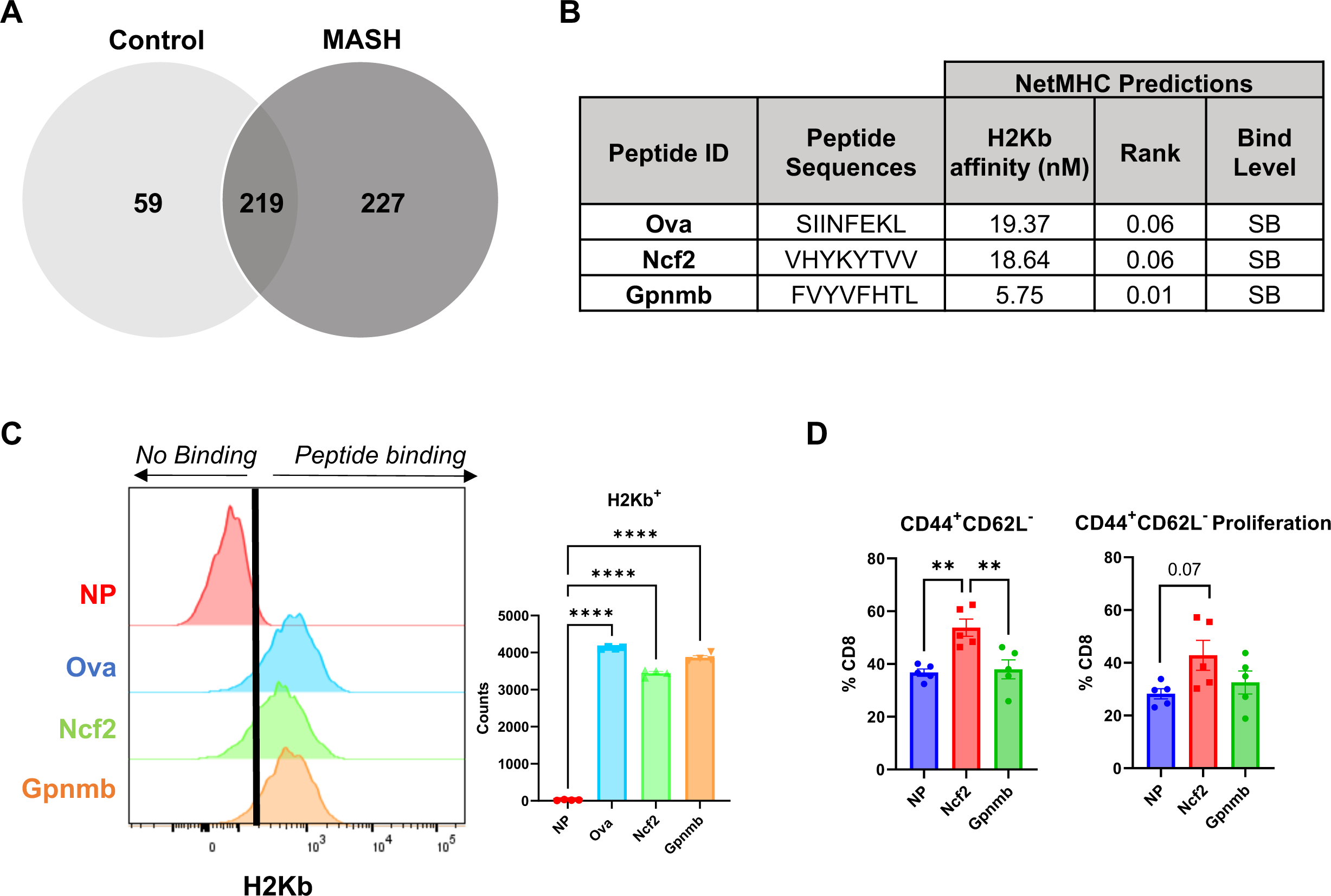
Identification of H2-K^b^ Peptides in MASH. LDLRKO mice on chow (control) or western diet (MASH) for 12 weeks. **(A)** BioVenn filtered unique control and MASH peptides. **(B)** NetMHC predicted binding affinities for Ovalbumin (Ova), Ncf2, and Gpnmb peptide to H2-K^b^. **(C)** Flow cytometry analysis of H2-K^b^ expression to determine binding with NP, Ova, Ncf2, and Gpnmb peptide pulsed RMA-S cells. **(D)** T cell activation assay of splenic CD8^+^ T cells co-cultured with NP, Ncf2, or Gpnmb peptide pulsed RMA-S cells. Cells were harvested on day 3 for flow cytometry analysis of CD8^+^ T cells (*CD8^+^TCRβ^+^)* and gated for activation (*CD44^+^ CD62L^-^*) and proliferation. Data shown as the mean ± SEM. Two-way ANOVA was performed and data were considered statistically significant for *P*<0.01 (**) and *P*<0.0001(****). *N* = 3–4 mice per group, 2 replicate studies.

To confirm H2-K^b^ peptide binding, a peptide binding assay was performed. Flow cytometry analysis confirmed H2-K^b^ binding for both the Ncf2 and Gpnmb peptides. The Gpnmb peptide also demonstrated a stronger affinity for H2-K^b^ compared with positive H2-K^b^ binding control Ovalbumin (Ova) (sequence: SIINFEKL) and Ncf2 peptide (Figure 1C). The Ncf2 peptide increased MASH spleen CD8^+^ T cell activation. In contrast, the Gpnmb peptide had no impact on the activation or proliferation of CD8^+^ T cell activation (Figure 1D).

The Gpnmb peptide was selected as a model peptide to initiate the development of our blocking peptides on the basis that the Gpnmb peptide has a high H2-K^b^ binding capability and is non-reactive towards MASH CD8^+^ T cells. The protein glycoprotein nonmetastatic melanoma protein B (Gpnmb) is an endogenous transmembrane glycoprotein that is upregulated during liver injury and fibrosis in mice[16]. Gpnmb undergoes cleavage and releases its extracellular domain into circulation[17, 18]. The extracellular domain can be detected in the serum and is upregulated in MASH mice, serving as a potential biomarker for MASH[19]. Interestingly, the identified peptide sequence for Gpnmb is in the extracellular domain region. Therefore, we did not select the Gpnmb peptide as a potential blocking peptide because it was identified in MASH livers and may have other functions in MASH development.

### Molecular modeling and computational identification of blocking peptides

*Pep*tide *B*inding *D*esign (PepBD) is a Monte-Carlo based search algorithm that uses an iterative procedure to identify high affinity peptide binders for a biomolecular target. The input to the algorithm is the structure of the complex formed by an initial peptide sequence (the “reference peptide”) and the target biomolecule. PepBD generates peptide variants by implementing sequence and conformational change moves on the peptide chain. A score function, *Γ_score_*, that measures the binding energy of the peptide to the receptor and the conformational stability of the peptide when bound to the receptor is used to accept or reject newly generated peptide sequences. Details of the algorithm are provided in the *Methods* and Supplementary material.

The reference peptide and input structure needed to initiate the PepBD search were determined by performing explicit-solvent atomistic molecular dynamics simulations of the Ncf2:H2-K^b^ and Gpnmb:H2-K^b^ peptide-protein complexes and evaluating their Δ*G_binding_*. To build the initial input files for performing molecular simulations, we obtained the coordinate file of peptide Ova bound to H2-K^b^ from the Protein Data Bank (PDB ID: 1VAC). The Ova peptide residues from the PDB file: 1VAC were mutated using the Pymol software to obtain the coordinate files of the Ncf2:H2-K^b^ (Supplementary Figure S1A) and Gpnmb:H2-K^b^ (Supplementary Figure S1B) complexes. Three independent simulations were carried out for both the Ncf2:H2-K^b^ and Gpnmb:H2-K^b^ complexes for 100 ns to ensure the system reached an equilibrated state. The simulations were performed at 298K using the AMBER ff14SB forcefield and the AMBER18 package. We calculated the Δ*G_binding_* of the peptide:receptor complex following the MD simulations using the MMGBSA protocol and variable dielectric constant method (See details in the *Methods* and Supplementary material). The Δ*G_binding_* of Ncf2:H2-K^b^ and Gpnb:H2-K^b^ were found to be −1.36 kcal/mol and −7.97 kcal/mol, respectively. These values confirmed our peptide binding assay results that Gpnmb peptide binds with a stronger affinity than Ncf2 to H2-K^b^. The representative snapshot from the last 5 ns MD simulation of the Gpnmb:H2-K^b^ complex obtained by performing a hierarchical clustering algorithm is shown in Figure 2A.

**Figure 2.**
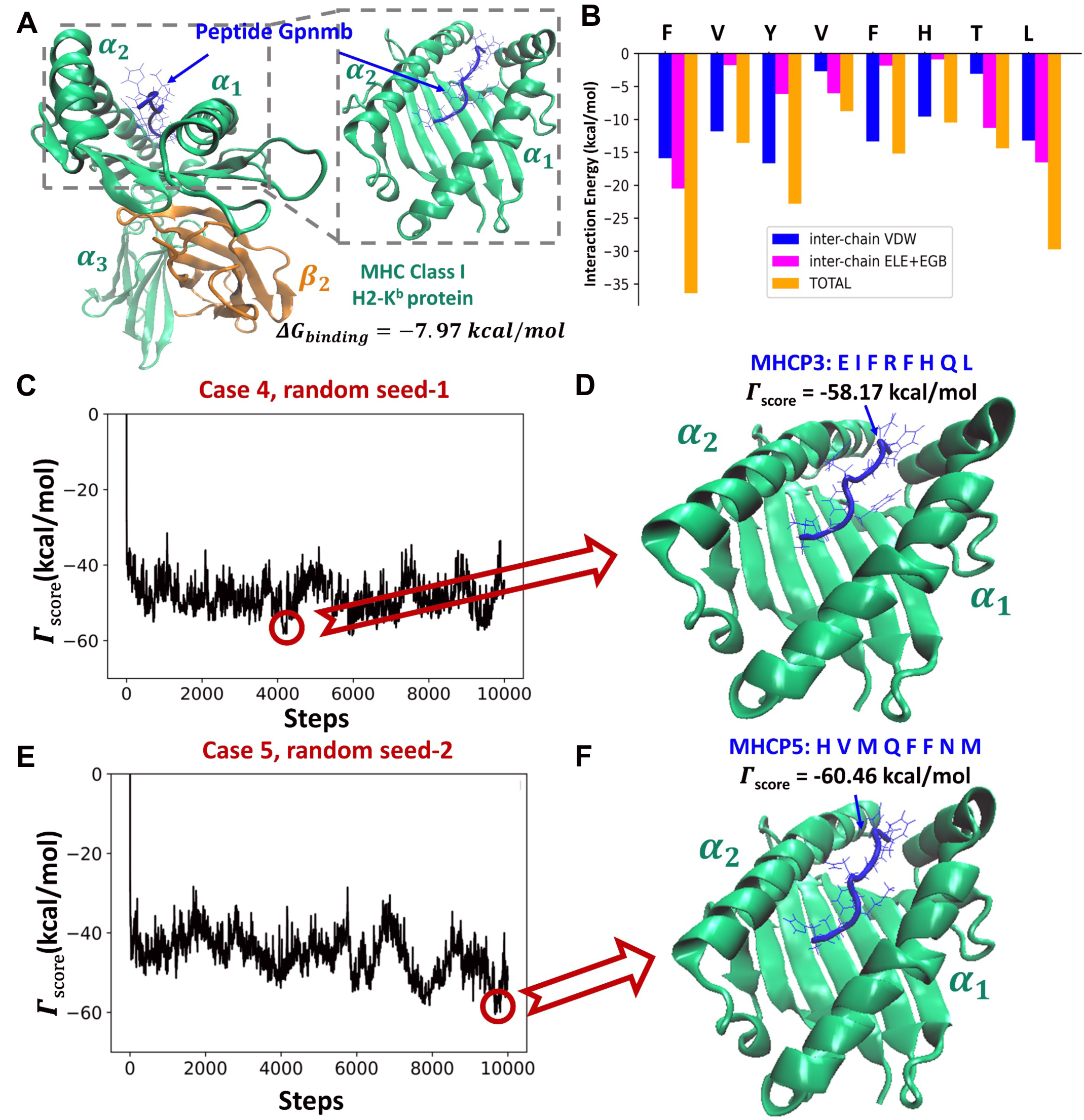
Molecular Interactions of Target and Blocking Peptides. **(A)** Peptide Gpnmb (FVYVFHTL) bound to MHC Class I H2-K^b^. The a_1_, a_2_, and a_3_ domains are shown in green. The {3_2_ domain is shown in orange. Peptide Gpnmb at the peptide-binding cleft of MHC Class I H2-K^b^ is enlarged to highlight the molecular interactions between the peptide and H2-K^b^. **(B)** The residue-wise interaction energy decomposition of Gpnmb when bound to H2-K^b^. The Γ*_score_* versus the number of steps for **(C)** Case 4 (random seed-1) resulting in **(D)** MHCP3:H2-K^b^ complex and for **(E)** Case 5 (random seed-2) resulting in **(F)** MHCP5:H2-K^b^ complex.

We selected peptide Gpnmb as the reference peptide and the Gpnmb:H2-K^b^ complex (Figure 2A) as the input structure for our PepBD algorithm. The goal was to generate new peptide sequences using PepBD that bind to H2-K^b^ with higher binding affinity than Gpnmb. The individual contributions from the residues on Gpnmb to the interaction energy with H2-K^b^ are plotted in Figure 2B. For our designs, we considered five different cases for PepBD that specify different amino acid composition by residue type for the peptide chain. Amino acids are classified into five types: hydrophobic, hydrophilic, positive charge, negative charge, and glycine. The numbers of each type are fixed during the design process to improve peptide stability and increase the likelihood the peptide will bind to the target. For example, the number of hydrophobic residues is fixed to enhance the likelihood the peptide will be soluble, and the number/position of charged amino acids is fixed to complement the charge state of the protein. The five cases are as follows. Case One: *N*_hydrophobic_ = 3, *N*_hydrophilic_ = 2, *N*_positive_ = 2, *N*_negative_ = 1, *N*_other_ = 0 and *N*_glycine_ = 0, Case Two: *N*_hydrophobic_ = 3, *N*_hydrophilic_ = 2, *N*_positive_ = 1, *N*_negative_ = 1, *N*_other_ = 0 and *N*_glycine_ = 1, Case Three: *N*_hydrophobic_= 3, *N*_hydrophilic_ = 2, *N*_negative_ = 2, *N*_positive_ = 1, *N*_negative_ = 2, *N*_other_ = 0 and *N*_glycine_ = 0, Case Four: *N*_hydrophobic_= 4, *N*_hydrophilic_ = 2, *N*_negative_ = 3, *N*_positive_ = 1, *N*_negative_ = 1, *N*_other_ = 0 and *N*_glycine_ = 0 and Case Five: *N*_hydrophobic_= 5, *N*_hydrophilic_ = 3, *N*_negative_ = 0, *N*_positive_ = 0, *N*_negative_ = 0, *N*_other_ = 0 and *N*_glycine_ = 0. (Here, *N*_residue-type_ stands for the number of residues specific to that residue type on the peptide chain). For each case, we performed the PepBD search with three different initial random seed numbers that randomize the initial peptide sequence. We do this to let PepBD, which is a Monte-Carlo search, sample peptides from a large pool of peptide sequences and conformations to avoid getting trapped in local minima. As the *in-silico* search proceeds and new peptide sequences and conformers are generated, the Γ*_score_*is recorded at each step. A lower Γ_score_ means stronger binding affinity of the peptide to the bound target. Figure 2C shows the Γ*_score_* vs the number of sequence and conformational change moves performed with random seed-1 for Case 4. Figure 2D shows the structure of one of the top performing peptides, MHCP3 (sequence: EIFRFHQL) complexed with H2-K^b^ from Case 5. Figure 2E shows the Γ*_score_* versus the number steps performed with random seed-2 for Case 5. The MHCP5: *HVMQFFNM* complexed with H2-K^b^, obtained from the corresponding search, is shown in Figure 2F.

Once the *in-silico* evolution of the peptide sequence is complete, we performed explicit-solvent atomistic MD simulations of the lowest scoring peptides complexed with the H2-K^b^ protein to predict their binding affinity, Δ*G_binding_*. We performed three 100 ns independent simulations for each peptide:H2-K^b^ complex. Each independent simulation starts with the same initial coordinates of the peptide:H2-K^b^ complex but with randomized velocities drawn from a Gaussian distribution. The Δ*G_binding_* of the peptide:receptor complex following the MD simulations was evaluated by using the MMGBSA protocol and variable dielectric constant method. Details of our atomistic MD simulation and Δ*G_binding_* calculation procedure are provided in the *Methods* and Supplementary material. Table 1 reports the top peptides obtained from our computational procedure and the corresponding Γ*_score_* and Δ*G_binding_* values.

**Table 1.**
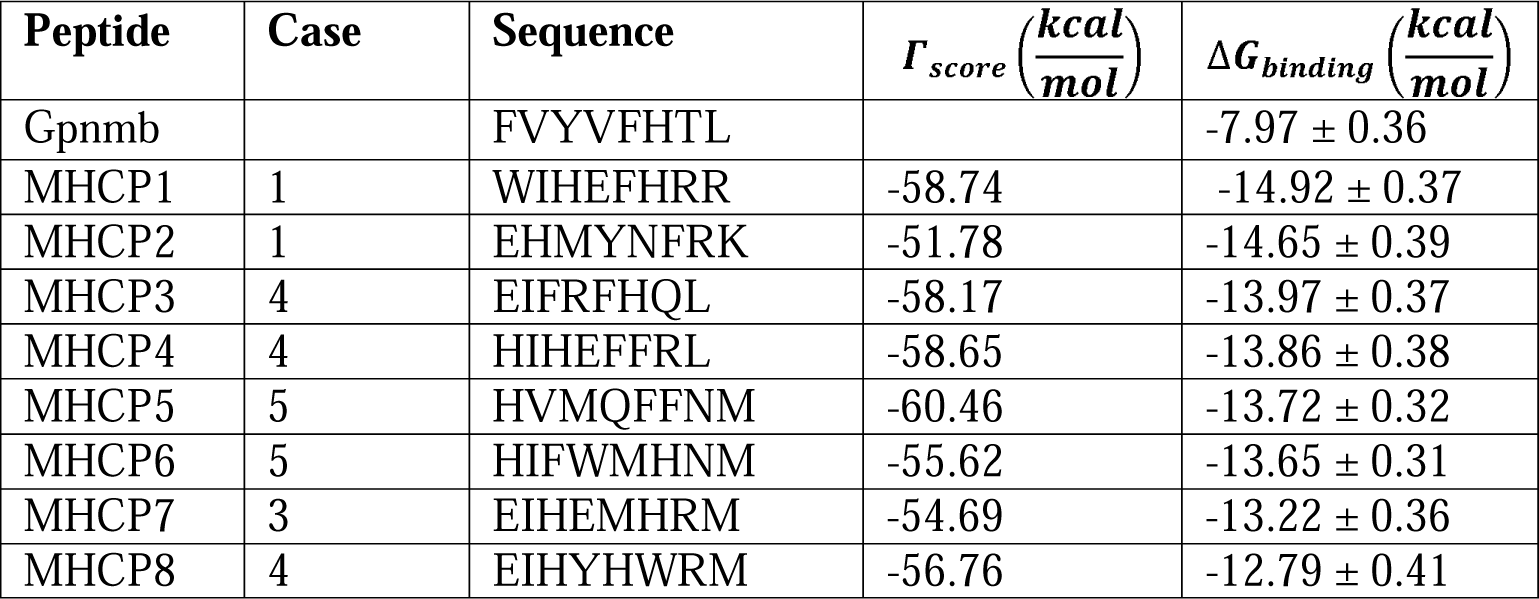
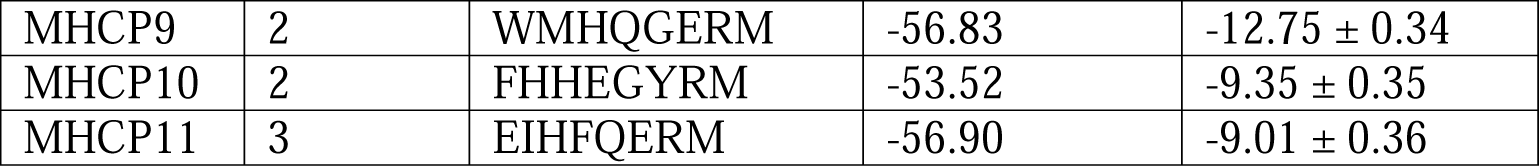
The initial peptide sequence (Gpnmb) and the list of peptide sequences identified by PepBD screening with the corresponding Δ*Γ_score_* and Δ*G_binding_* values.

### Computational analysis of the biorecognition mechanism of MCHP3 and MHCP5 bound to H2-K^b^

Next, we analyzed the ability of two of the designed peptides, MHCP3 and MHCP5, to bind to H2-K^b^ with a particular focus on which amino acids were key to the binding. The focus was on peptides MHPC3 and MHCP5 because, as we show in the next sections, these featured moderate-to-high binding affinity for H2-K^b^ and protected against Ncf2 CD8^+^ T cell activation. The Δ*G_binding_* of MHCP3:H2-K^b^ (Figure 3A) and MHCP5:H2-K^b^ (Figure 3B) from atomistic molecular dynamics simulation were predicted to be −13.97 kcal/mol and −13.72 kcal/mol, respectively. To draw further insights on the binding mechanism of MHPC3 and MHCP5 to H2-K^b^, we constructed the residue-wise decomposition of the interaction energy between MHCP3:H2-K^b^ (Figure 3C) and MHCP5:H2-K^b^ (Figure 3D) binding interface. In Figures 3E and 3F, we also constructed energy panels detailing the pair-wise interactions of MHCP3:H2-K^b^ and MHCP5:H2-K^b^ complexes, respectively.

**Figure 3.**
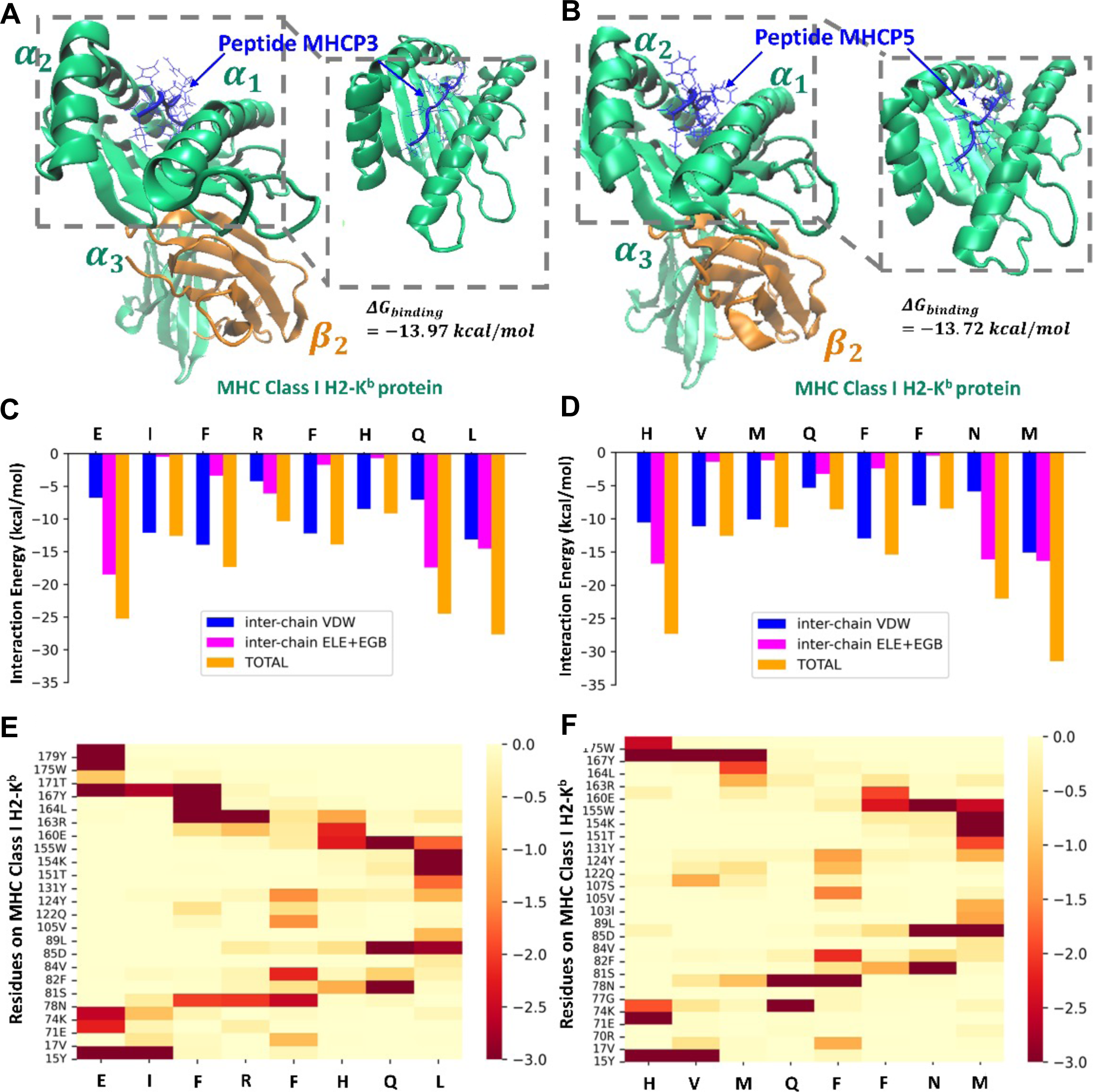
Molecular Structure of MHCP3 and MHCP5 Binding to H2-K^b^. The structure of **(A)** MHCP3:H2-K^b^ and **(B)** MHCP5:H2-K^b^ obtained from molecular dynamics simulations. The residue-wise interaction energy decomposition of **(C)** MHCP3:H2-K^b^ and (D) MHCP5:H2-K^b^ complexes. Energy panel detailing the pair-wise interactions in **(E)** MHCP3:H2-K^b^ and **(F)** MHCP5:H2-K^b^ complexes.

The critical MHCP3 residues involved in H2-K^b^ binding are Glu1, Phe3, Gln7, and Leu8, while the critical residues on MHCP5 are His1, Asn7, and Met8. Glu1 on MHCP3 interacts strongly with Tyr15, Tyr167, Trp175, and Tyr179 on H2-K^b^. Because Glu1 (negatively charged) was forming strong interactions with Tyr and Trp, which are both aromatic amino acids, we suspect these interactions are via anion-π contacts. Phe3 on MHCP3 preferentially interacts with Arg163, Leu164, and Tyr167, possibly via cation-π, Van der Waals contacts and π-π interaction, respectively. Gln7 (polar) on MHCP3 interacts with polar Ser81 and Asp85 on H2-K^b^ via hydrogen bonds and with Trp155 via polar-π interaction. Leu8 on MHCP3 forms strong interactions with Asp85, Thr151, and Lys154. His1 on MHCP5 interacts with Tyr15 and Tyr167 via polar-π interactions and with Glu71 via hydrogen bonding. Asn7 on MHCP5 interacts strongly with Ser81 and Asp85 on H2-K^b^ via hydrogen bonding and with Trp155 on H2-K^b^ via polar-π interaction. Met8 on MHCP5 interacts with Asp85, Thr151, and Lys154. Notably, both MHCP3 and MHCP5 contain two Phe (large aromatic amino acid) residues and may provide conformational stability to the peptide chain.

### Experimental Evaluation of the Ability of Blocking Peptides MHCP3 and MHCP5 to bind to H2-K^b^

We synthesized blocking peptides MHCP1 to MHCP11 and determined the ability to bind to H2-K^b^ by using NetMHC predictions (Figure 4A) and H2-K^b^ peptide binding by flow cytometry analysis (Figure 4B). MHCP 3, 4, 5, and 6 were all predicted to be strong binders by NetMHC. By pulsing RMA/S cells with the blocking peptides and detecting stabilization of H2-K^b^ on the cell surface via flow analysis, the blocking peptides MHCP3 and MHCP5 bound to H2-K^b^. The binding of MHCP4 to H2-K^b^ was not as strong as that experienced by MHCP3 and MHCP5. Interestingly, the presence of two Phe in the blocking peptides plays an important role in the binding strength of the blocking peptides as peptides MHCP1, 2, 6, 7,8, 9, 10, and 11 did not bind to H2-K^b^.

**Figure 4.**
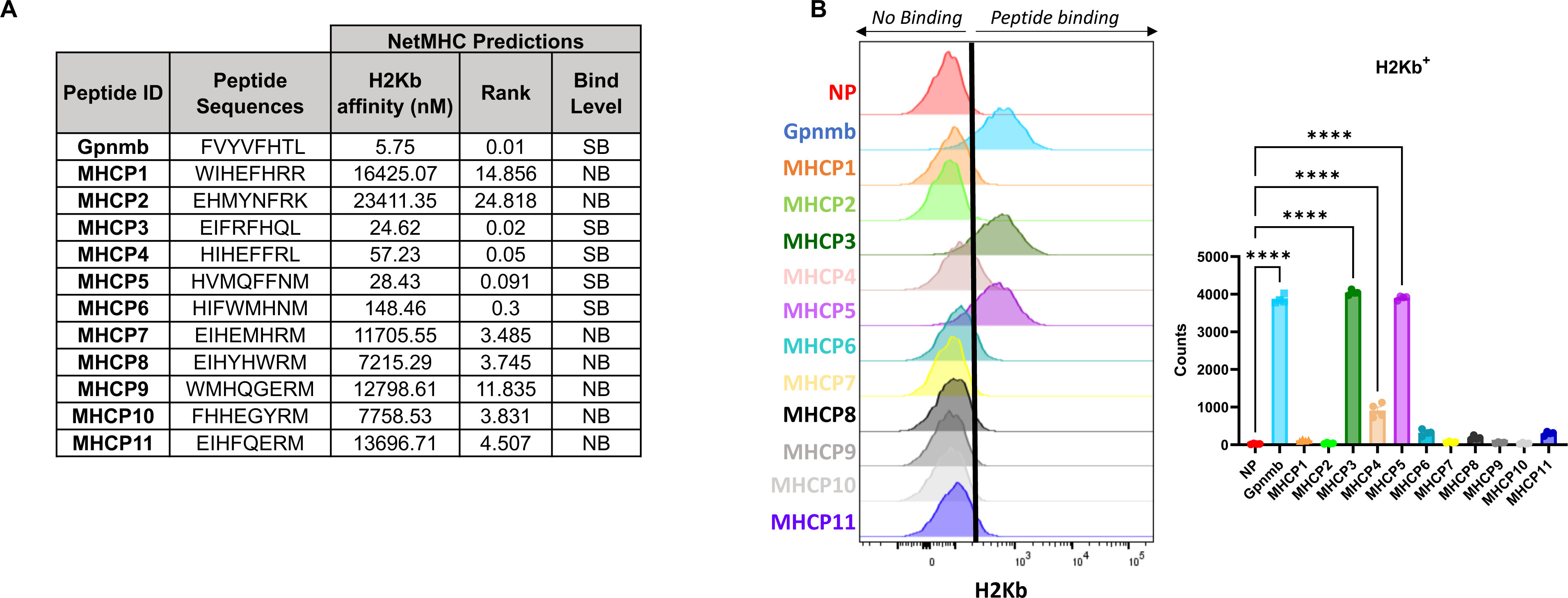
MHCP3 and MHCP5 bind to H2-K^b^. **(A)** NetMHC predicted binding affinities for generated blocking peptides compared with Gpnmb peptide. **(B)** Flow cytometry analysis of H2-K^b^ expression to determine binding with blocking peptides (*n*=6, in 3 replicate studies). Data shown as the mean ± SEM. Two-way ANOVA was performed and data were considered statistically significant for *P*<0.0001(****). *N*=3 and 2 replicate studies.

### Blocking peptides MHCP3 and MHCP5 protect against Ncf2 CD8^+^ T cell activation

To determine the ability of the blocking peptides to prevent Ncf2-induced CD8^+^ T cell activation, RMA-S cells were treated with blocking peptides MHCP3 or MHCP5 and the Ncf2 peptide. RMA-S cells were either pretreated with MHCP3 or MHCP5 for 1 hour (PreT) and then loaded with Ncf2 peptide or simultaneously treated with MHCP3 or MHCP5 and Ncf2 peptide for 5 hours (CoT). RMA-S cells were then cultured with isolated MASH splenic CD8^+^ T cells and analyzed for CD8^+^ T cell activation and proliferation. Both blocking peptides showed no CD8^+^ T cell reactivity on their own. Additionally, both MHCP3 and MHCP5 were able to prevent Ncf2-induced CD8^+^ T cell activation regardless of being pre-treated or co-treated with the Ncf2 peptide (**Figures 5A-B, 6A-B**). Our findings demonstrate that MHCP3 and MHCP5 are competitively effective in preventing Ncf2 peptide-induced activation of CD8^+^ T cells *in vitro*.

**Figure 5.**
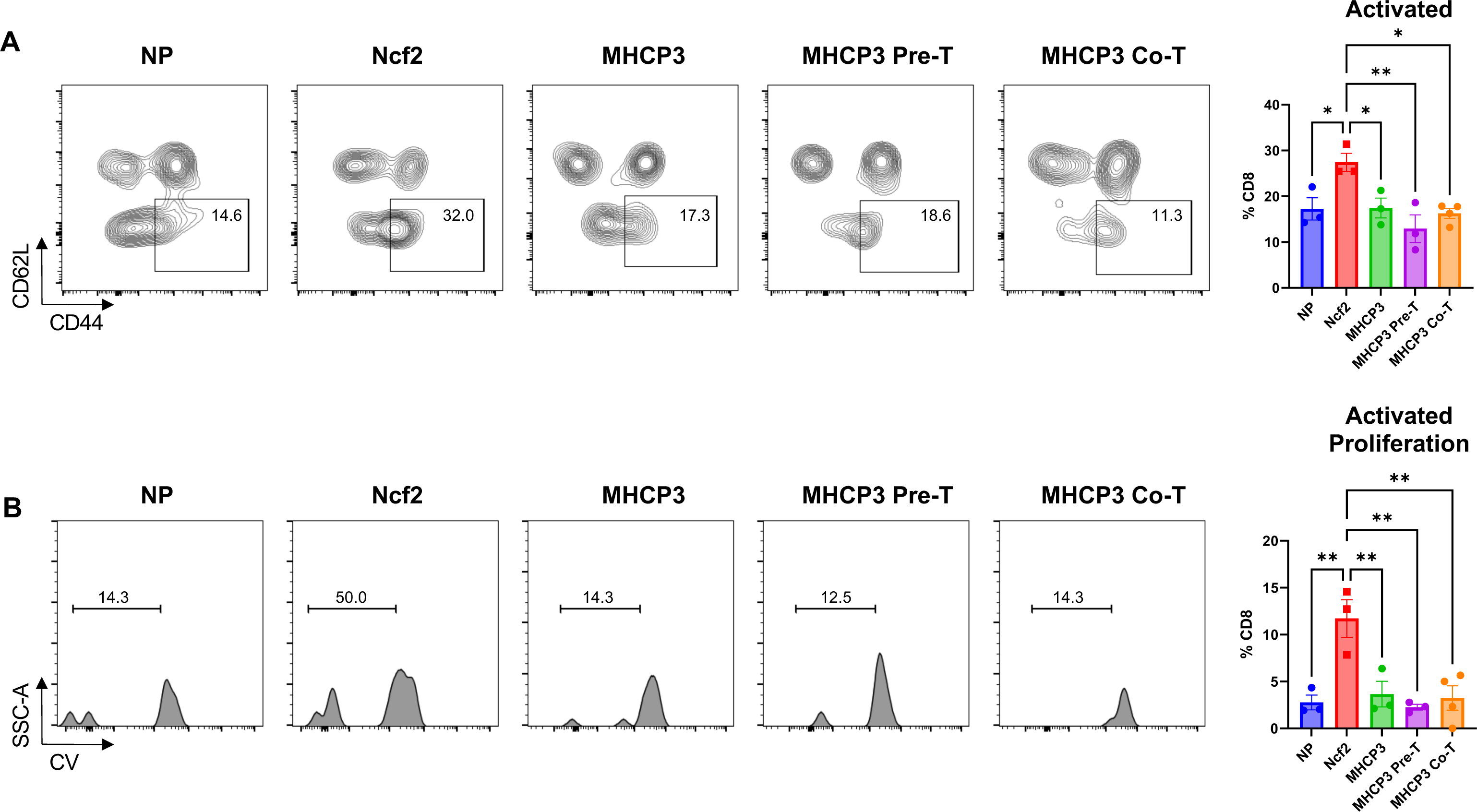
MHCP3 blocks Ncf2 peptide induced CD8^+^ T cell function. RMA-S cells were treated with no peptide (NP) or 100 µM Ncf2 peptide or MHCP3. RMA-S were pre-treated with MHCP3 (Pre-T) or treated in combination (Co-T) with the Ncf2 peptide. RMA-S cells were cultured with isolated MASH splenic CD8^+^ T cells for 5 days. Cells were harvested for flow cytometry for analysis of **(A)** CD8^+^ T cell activation (*CD44^+^, CD62L^-^*) and **(B)** proliferation. Data shown as the mean ± SEM. Two-way ANOVA was performed and data were considered statistically significant for *P*<0.05 (*), *P*<0.01 (**), *P*<0.001(***), and *P*<0.0001(****). *N*=3 and 2 replicate studies.

**Figure 6.**
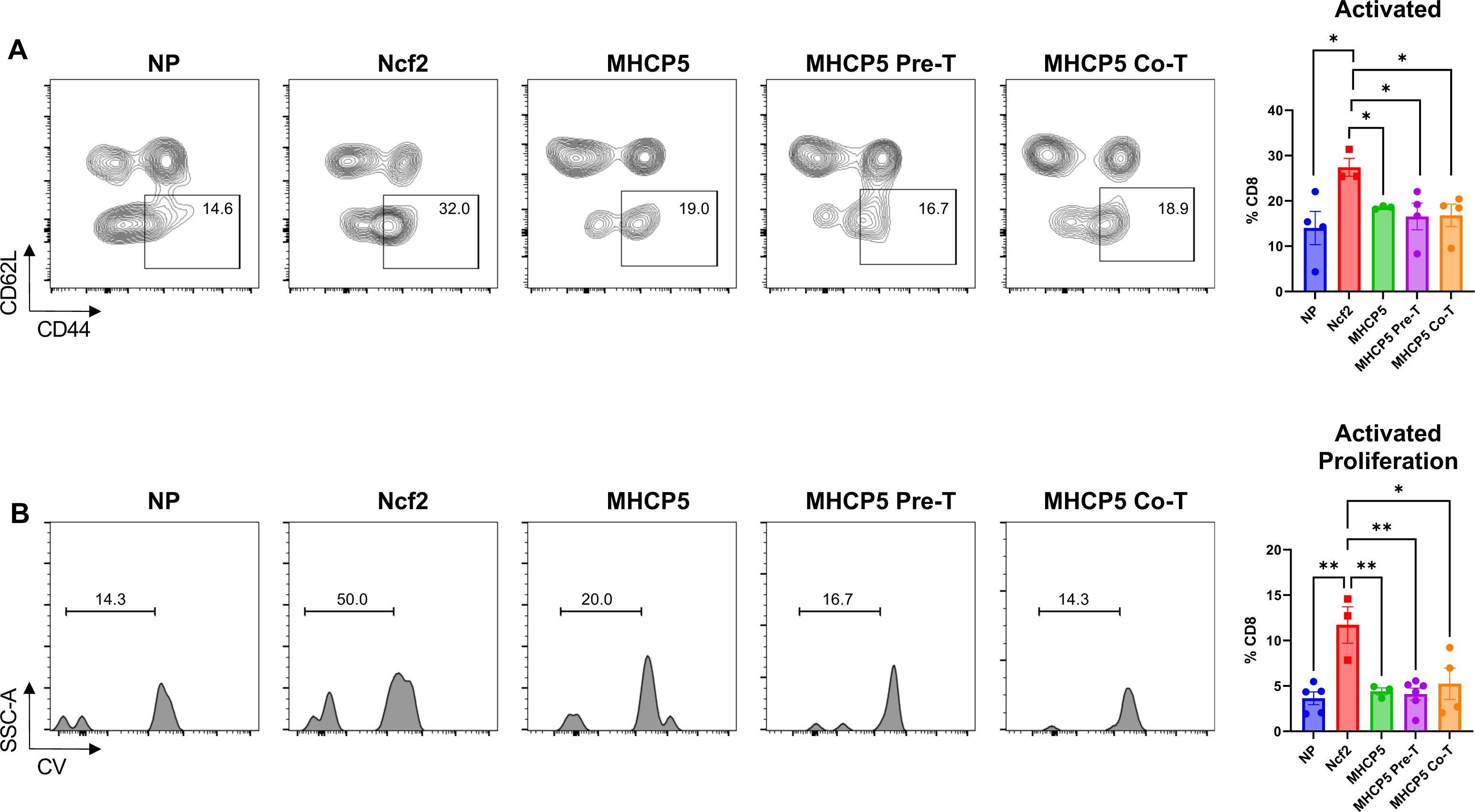
MHCP5 blocks Ncf2 peptide induced CD8^+^ T cell function. RMA-S cells were treated with no peptide (NP) or 100 µM Ncf2 peptide or MHCP5. RMA-S cells were pre-treated with MHCP5 (Pre-T) or treated in combination (Co-T) with the Ncf2 peptide. RMA-S cells were cultured with isolated MASH splenic CD8^+^ T cells for 5 days. Cells were harvested for flow cytometry for analysis of **(A)** CD8^+^ T cell activation (*CD44^+^, CD62L^-^*) and **(B)** proliferation. Data shown as the mean ± SEM. Two-way ANOVA was performed and data were considered statistically significant for *P*<0.05 (*), *P*<0.01 (**), *P*<0.001(***), and *P*<0.0001(****). *N*=3 and 2 replicate studies.

## DISCUSSION

CD8^+^ T cells play a critical role in the development of MASH. Identifying MASH-specific immunopeptides that regulate CD8^+^ T cell activation has opened new possibilities for immunotherapies targeting specific antigens bound to MHC class I. Blocking peptides may be better than antibody therapy due to the smaller size and ease of altering stability and specificity. Developing competitive binding blocking peptides with high binding affinity to MHC class I molecules H2-K^b^ may prevent CD8^+^ T cell activation and serve as a possible treatment for MASH. We discovered two synthetic blocking peptides containing 8 amino acids that specifically interact with H2-K^b^ using the PepBD algorithm, which searches for peptides with high affinity and selectivity to a protein target. Both blocking peptides have a strong binding affinity to H2-K^b^ and competitively inhibit antigen activation of MASH CD8^+^ T cells.

Computational design strategies are beneficial for developing targeted blocking peptides that bind to the target of interest with higher specificity and efficacy than a reference peptide. By using the Gpnmb peptide as a reference peptide due to its strong binding affinity and the PepBD algorithm coupled with molecular-level simulations, we were able to design peptides with stronger binding affinity to H-2K^b^ than the Ncf2 and Gpnmb peptides. Interestingly, NetMHC 4.0 predicted MHCP3 and MHCP5 to be strong binders. In addition, both blocking peptides demonstrated stable binding to the H-2K^b^ receptor on RMA-S cells *in vitro* compared with the other predicted blocking peptides. The stable binding may be due to the presence of critical amino acids in the MHCP3 and MHCP5, as both contain Phe. Future studies mutating Phe in the blocking peptides may help determine the functional significance of this amino acid and further the development of stable and effective blocking peptides.

Several groups have demonstrated the stability of peptides to the MHC I and II molecules controls immunogenicity versus affinity[20–33]. We discovered that the blocking peptides did not induce an immunogenic response in hepatic isolated T cells. In addition, we analyzed if the blocking peptide sequences were present in the immunopeptidome of normal, steatosis, or MASH mouse livers and found these were not present under such conditions. This finding further supports the possibility that these blocking peptides may be effective *in vivo* for treating MASH. Future *in vivo* testing will confirm the immunogenic capacity of MHCP3 and MHCP5.

Delivery of peptides can be challenging as certain amino acids lead to poor bioavailability, membrane permeability, and short half-life. Modifications to enhance protease stability by incorporating D-amino acid or adding acetyl or propionyl groups to the N- and C-terminal amino acids may improve the delivery and effectiveness of the blocking peptides *in vivo*. Exosomes or nanoparticles are a possible peptide delivery method. We have demonstrated that myeloid cells are a critical antigen-presenting cell in regulating CD8^+^ T cell activation in MASH (Adams et al.). Thus, developing exosomes containing MHCP3 and MHCP5 may effectively target antigen-presenting cells, preventing CD8^+^ T cell activation. These exosomes may be delivered by systemic administration of macrophage-derived exosomes, encapsulated exosomes in carriers such as nanoparticles, or modification of the membrane of exosomes to enhance binding to receptors on antigen-presenting cells. With the lack of FDA-approved MASH therapies, the blocking peptides we designed may be an effective treatment for MASH and other metabolic diseases dependent on antigen presentation and T cell function.

## MATERIALS AND METHODS

### Animal Models

*LDLRKO MASH model*. 5-week-old male low-density lipoprotein receptor knockout (LDLRKO) mice were originally purchased from Jackson Laboratories (Bar Harbor, ME) and propagated further in our colony. 6-week-old male LDLRKO mice were fed western diet (WD, 42 Kcal% fat with 0.2% added cholesterol, TD.22137; Harlan Laboratories) for 12 weeks. All animal procedures were approved by the Institutional Animal Care and Use Committee (IACUC) at North Carolina State University under protocol 21-502-B.

### Isolation and identification of H2-K^b^ Peptides from Liver

Peptides were isolated and identified using methods previously described (Adams et al. submitted).

### Computational Peptide Design

*Pep*tide *B*inding *D*esign (PepBD) is a Monte-Carlo search algorithm that identifies peptide sequences that bind with higher affinity and specificity to a biomolecular target than a known “reference ligand”[11, 14, 34, 35]. The starting input structure for the algorithm is a complex formed between the reference ligand (in this case Gpnmb) and the target protein of interest (in this case H2-K^b^). The design algorithm iterates through 10,000 evolution steps and generates variants to the original peptide that bind to H2-K^b^. Two kinds of moves, sequence change and conformational change moves, are performed to generate peptide variants. After each move, the score (a measure of the peptide-target binding energy) for each trial peptide is compared with that of the previous sequence/conformation and accepted or rejected using the Monte-Carlo Metropolis sampling technique. The best peptide variants, meaning those with the best “scores,” are evaluated in explicit-solvent atomistic molecular dynamics simulations to calculate the binding free energy, Δ*G_binding_*, of the peptide:H2-K^b^ complex.

### Atomistic Molecular Dynamics Simulation

Explicit-solvent atomistic MD simulations were carried out in the canonical (NVT) ensemble using the AMBER 18 package to examine the dynamics of the binding process between the peptide:H2-K^b^ complexes. We completed three independent simulations for each peptide:H2-K^b^ complex. Each simulation started with a 1000 step solvent minimization with restraint force = 500 kcal/mol. This was followed by a 2500 step unconstrained energy minimization. The production simulations were performed for 100 ns. We solvated each peptide-receptor complex in a periodically truncated octahedral box containing a 12 Å buffer of TIP3P water (∼ 22,700 water molecules) surrounding the complex in each direction. Particle Mesh Ewald (PME) summation was used to calculate the long-ranged electrostatic interactions with a cut-off radius of 8 Å and a 1 × 10^-5^ tolerance for the Ewald convergence to calculate the nonbonded interactions. Hierarchical clustering analysis was performed on the last 5 ns of the simulation trajectories to obtain the representative structure of the peptide:H2-K^b^ complex. The last 5 ns simulation trajectory of each peptide:H2-K^b^ complex was evaluated to calculate the binding free energy by using the implicit-solvent molecular mechanics/generalized Born surface area (MM/GBSA) approach with the variable internal dielectric constant model. More details regarding our computational procedures and post-analysis of the atomistic MD simulations can be found in our previous work[14, 34, 36, 37].

### Peptide Synthesis

All peptides were synthesized at a crude purity from Peptide 2.0, Inc (Chantilly, VA). Synthetic peptides were used for peptide binding and T cell activation assays.

### H2-K^b^ Peptide Binding Assay

RMA-S cells were seeded into a 96-well plate at 1×10^5^ per well in 100 µL of media (RPMI with 10% FBS) and incubated at 27°C for 18 hours as described previously (Ncf2 ref). Following incubation cells were treated with 100 µM for each of the following peptides: no peptide vehicle control (NP), Ncf2 peptide, MHCP3, or MHCP5. Additionally, blocking peptides MHCP3 and MHCP5 were treated in combination with the Ncf2 peptide to evaluate blocking potential. All peptide treatments were incubated at 37°C for 5 hours and harvested for flow cytometry. All samples were incubated with Fc block for 5 minutes on ice, followed by incubation with fluorophore conjugated antibody H2-K^b^ (APC-eFluor780, 1:200; ThermoFisher) for 30 minutes. Cells were then stained with propidium iodide (PI, 1:10,000; ThermoFisher). Data for this assay were acquired on a Becton Dickinson LSRII machine in the NCSU Flow Cytometry Core and analyzed using FlowJo software v10.8. Flow cytometry gating strategy is displayed in Supplemental Figure 2A.

### CD8^+^ T cell Isolation from Spleen

Mouse spleens were strained though 100-µm filters with FACS buffer and pelleted. Pellets were washed with FACS buffer and pelleted again. Pellets were then incubated with ACK lysing buffer on ice for 5 mins after which FACS buffer was added and cells pelleted. Pellets were then washed twice and strained through a cell strainer cap and prepared for T cell activation assays.

### CD8^+^ T Cell Activation Assay

Isolated CD8^+^ T cells from MASH spleens were plated in 96-well plates at 5×10^4^ per well and stained with a cell trace violet proliferation dye and cocultured with 1×10^4^ of peptide treated RMA-S cells as described above. Cells were plated in a total of 200 µL of T cell media containing RPMI medium (Corning) supplemented with 10% FBS, L-glutamine (400 mM), penicillin (100 U/ml), streptomycin (100 µg/ml), 2-mercaptoethanol (50 µM), and IL-2 (100 ng). Cells were cultured at 37°C and harvested after 5 days and prepared for flow cytometry. Cells were washed twice with FACS buffer and incubated with Fc block before staining. Cells were stained with the following panel: CD8a (PE-Cy7, 1:200; BD Biosciences), TCRb (APC-Cy7, 1:200; BD Biosciences), CD44 (A700, 1:200; BD Biosciences), CD62L (APC, 1:200; BD Biosciences), and propidium iodide (PI). Flow data were obtained on the Beckman Coulter CytoFLEX machine at the NCSU Flow Cytometry Core and analyzed using FlowJo software v10.8. Flow cytometry gating strategy is displayed in Supplemental Figure 2B.

### Statistical Analysis

All statistical analyses were performed using GraphPad Prism 9.3.1 software. All graphs are displayed as the mean ± SEM. Two-way ANOVA was performed and considered statistically significant for P<0.05 (*), P<0.01 (**), P<0.001(***), and P<0.0001(****).

## DATA AVAILABILITY STATEMENT

The datasets presented in this study will be made available upon request to the corresponding author.

## ACKNOWLEDGEMENTS

This research was supported by funds from the National Science Foundation, CBET 1934284 and OAC 1931430 to CH. Flow Cytometry experiments were performed in the Flow Cytometry and Cell Sorting facility at North Carolina State University, College of Veterinary Medicine.

## AUTHOR CONTRIBUTIONS

VRA conducted *in vitro* and *in vivo* experiments and data curation. SS conducted computational design and data curation. CH and AK conceived and designed the studies. VRA and SS wrote the first draft of the manuscript with guidance from CH and AK. VRA, SS, CH and AK edited the manuscript.

