## Supplemental Figures 1-2 for "Development of MHC Class I Blocking Peptides to Target Metabolic Dysfunction-Associated Steatohepatitis CD8^+^ T Cell Activation"

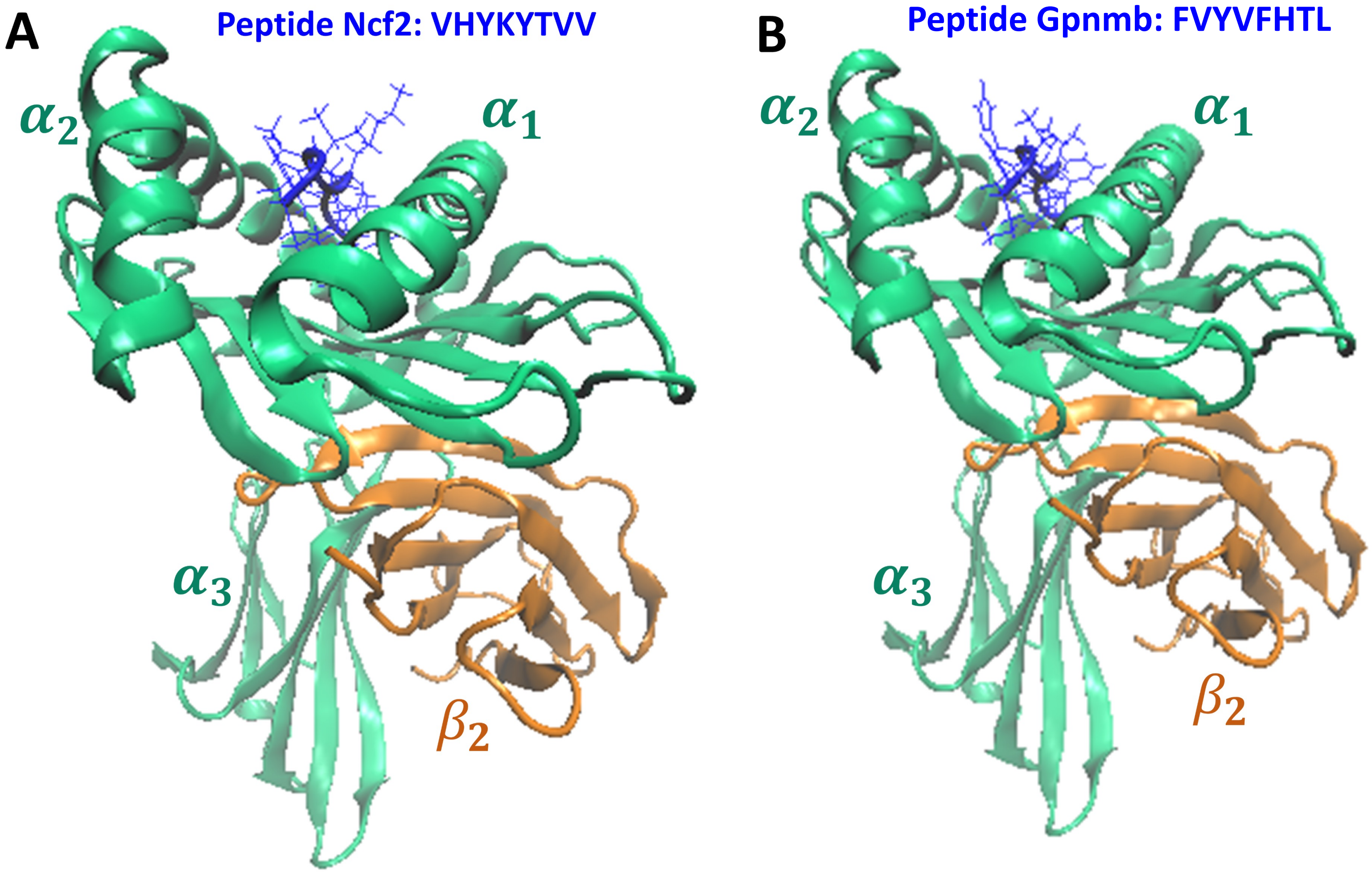


**Figure S1. (A)** Peptide Ncf2 (VHYKYTVV) bound to MHC Class I H2-Kb. The $\alpha_1$, $\alpha_2$ and $\alpha_3$ domains are shown in green. The $\beta_2$ domain is shown in orange. Peptide Ncf2 at the peptide-binding cleft of MHC Class I H2-Kb is enlarged to highlight the molecular interactions between the peptide and H2-Kb. **(B)** Peptide Gpnmb (FVYVFHTL) bound to MHC Class I H2-Kb. The $\alpha_1$, $\alpha_2$ and $\alpha_3$ domains are shown in green. The $\beta_2$ domain is shown in orange. Peptide Gpnmb at the peptide-binding cleft of MHC Class I H2-Kb is enlarged to highlight the molecular interactions between the peptide and H2-Kb.

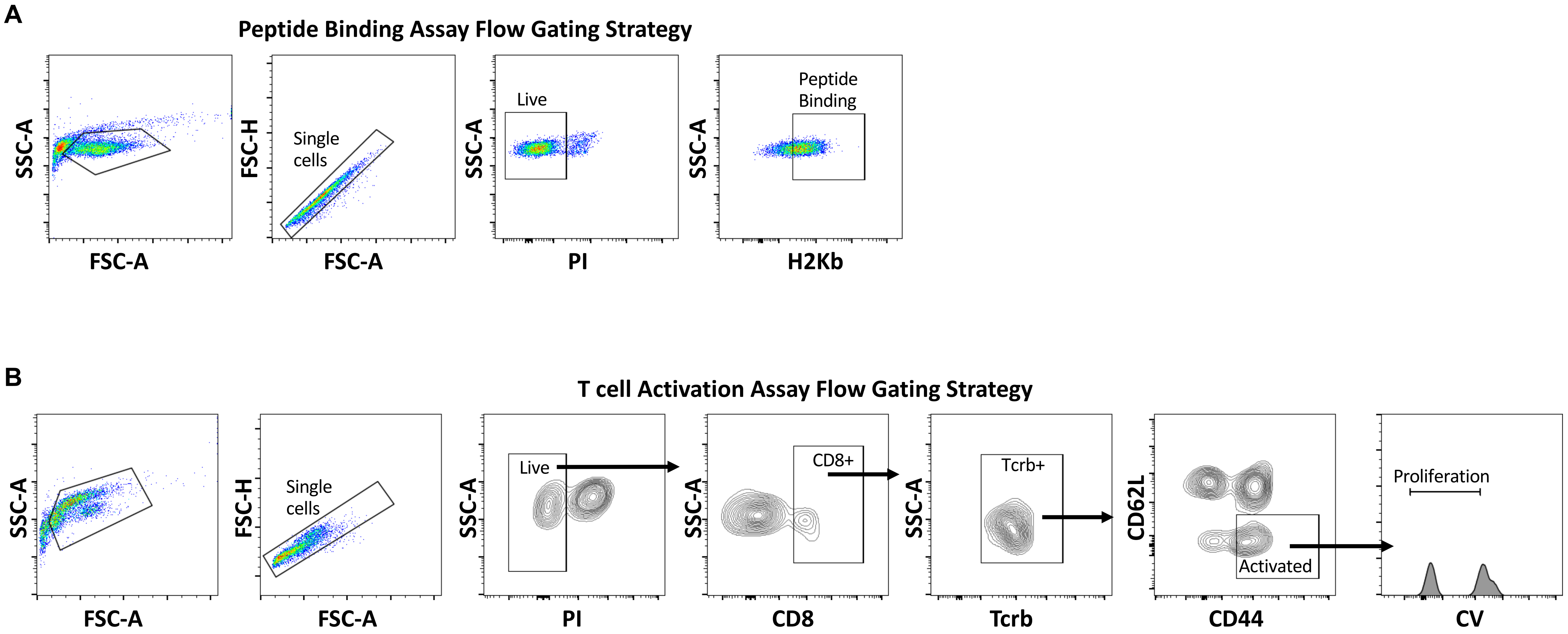


**Figure S2. (A)** Flow cytometry T cell activation gating strategy. **(B)** Representative flow plots for T cell activation assay with RMA-s pulsed with NP, Ncf2, or Gpnmb peptides. **(C)** Flow gating strategy for peptide binding assay. Data shown as the mean ± SEM. Two-way ANOVA was performed and considered statistically significant for P<0.01 (**).
